# Ontology-Driven Annotation of Key Events Enables Transcriptomics-Based Assessment of Adverse Outcome Pathway Networks

**DOI:** 10.64898/2026.10.06.756965

**Authors:** Sidra Adil, Theo M. de Kok, Egon L. Willighagen, Tooba Abbassi-Daloii, Marvin Martens

## Abstract

Chemical safety assessment is moving towards new approach methodologies (NAMs) to improve human safety and reduce animal testing. The Adverse Outcome Pathway (AOP) is a NAM framework describing toxicity as a chain of Key Events (KEs). Many AOP-Wiki KEs describe molecular, cellular, or tissue-level processes and can therefore be assessed with gene expression data, making transcriptomics of human *in vitro* models a NAM data generator. In AOP-Wiki, KEs are described by Key Event Components (KECs). A gene set can be derived when the biological process (BP) component carries a Gene Ontology (GO) term. Most KEs carry no such term, so their perturbation cannot be assessed using omics data. We assigned GO BP terms to the unmapped BP component across the brain, lung, liver and kidney AOP networks. We also replaced obsolete and direction-carrying terms in the pre-existing mapping with current, direction-neutral ones. For liver, we reviewed all pre-existing terms and replaced overly broad terms with better-matched alternatives. We then applied this approach to pesticide-induced hepatotoxicity, using Open TG-GATEs transcriptomics data from primary human hepatocytes exposed to cycloheximide, 2,4-dinitrophenol, hexachlorobenzene, and rotenone. KE perturbation was scored using single-sample Gene Set Enrichment Analysis (ssGSEA). Annotation increased the proportion of transcriptomics-linkable KEs in the liver network from 53% to 77%, enhancing identification of perturbed KEs after exposure. The KEs perturbed by all four compounds fell into five functional categories associated with hepatotoxicity. Our approach resulted in a more relevant AOP network for transcriptomics-based assessment of KE perturbation. These findings show molecular perturbation rather than direct proof of KE activation or causal progression through an AOP. When validated on larger and independent datasets, such networks will support mechanistic interpretation of transcriptomics data in chemical safety assessment.

## 1 Introduction

Predicting chemical toxicity requires understanding not only which biological processes a chemical disturbs, but how these disturbances translate into adverse effects. Adverse Outcome Pathways (AOPs) provide this mechanistic bridge, offering structured frameworks that describe cause-effect relationships between key events (KEs) across biological scales. Each AOP begins with a molecular initiating event (MIE), the first interaction between a chemical stressor and a biological target such as a receptor, enzyme, or membrane, followed by downstream KEs at the molecular, cellular and tissue levels that progressively lead to adverse outcomes (AOs) at organ, individual, or population levels [1, 2]. AOPs are increasingly used in toxicology and regulatory decision-making [3, 4]. Because KEs are modular [3], the same KE often appears in several AOPs. The AOPs that converge on damage to one organ therefore form a network of KEs rather than a set of independent chains. Making AOPs quantitative matters for chemical risk assessment, which requires mechanistic descriptions to be connected to measurable biological responses. While qualitative AOPs define the causal chain from MIE to AO, quantitative AOPs offer measurable descriptors for KEs and key event relationships (KERs), enabling predictive modelling [5, 6]. Qualitative AOPs can provide a basis for more quantitative applications when KEs are linked to measurable molecular data, such as gene expression profiles [7, 1, 8]. Quantitative AOPs are therefore central to new approach methodologies (NAMs), which are currently a priority in Europe, driven by the goal of reducing and replacing animal testing for ethical and animal-welfare reasons [9]. The case for NAMs is not only ethical. Animal tests often fail to predict what a chemical does in humans. Animal and human results agree for more than 80% of haematological, gastrointestinal, and cardiovascular effects, but for fewer than 35% of cutaneous ones [10]. Across a wider set of comparisons, animal and human studies agreed about half the time [11]. A systematic review found that these agreement rates vary widely between studies [12], and recent work points to limited reproducibility of the animal tests themselves [13]. Test systems built from human cells avoid the need to translate between species. In regulatory toxicology, the question is now how and when such methods are adopted, rather than whether [14].

For AOPs to be integrated with quantitative molecular data, their biological descriptions need to be standardised and computationally accessible, which are requirements met through ontological annotation. AOP-Wiki (https://aopwiki.org/) serves as the primary repository for AOPs, storing KEs as biological events that are measurable and essential to progression to AO. KEs are described by three key event components (KECs): biological process (BP), biological object (BO), and biological action (BA). BP defines what is happening, BO specifies the affected entity, and BA indicates directionality (e.g., increase/decrease). For instance, the KE 1115 ‘Increase, Reactive oxygen species’ is described by the BP ‘reactive oxygen species biosynthetic process’ (GO:1903409), the BO ‘reactive oxygen species’, and the BA ‘increased’. These KECs are standardised using ontologies, which are controlled vocabularies that assign machine-readable identifiers to biological entities and processes [15]. Ontologies ensure consistent interpretation across databases and enable precise KE descriptions by linking them to well-defined biological processes and objects [16], making AOP information findable, accessible, interoperable, and reusable (FAIR) [17]. In AOP-Wiki, BP and BO are described using existing ontologies (primarily Gene Ontology (GO) for BP, ChEBI and Protein Ontology for BO), while BA uses internal ontologies for directionality [16, 18]. This division matters for annotation. KE titles often contain a direction word, and KE 1115 begins with ‘Increase’. That direction is already recorded in the BA, so it should not influence the choice of BP term. A GO BP term that itself specifies a direction would duplicate the BA and return the genes that regulate a process rather than those that carry it out. BP annotation should therefore describe the process alone.

Despite this ontological framework, a substantial proportion of KEs in AOP-Wiki carry no BP description [18], so no gene set can be derived from them, and their perturbation cannot be measured with omics approaches. The annotations that are present are not always usable either. GO terms can become obsolete, some carry a direction rather than describing the underlying process, and others sit at a level of the ontology so general that the resulting gene set extends far beyond the event the KE describes. These gaps matter because most KEs represent molecular and cellular biological processes suitable for transcriptomics-based assessment, where gene expression changes can reflect KE perturbation. A GO BP annotation is what allows a gene set to be derived for a KE. Where that annotation is missing or poorly fitting, no usable gene set can be assigned, and the KE cannot be assessed with transcriptomics data. This limits the integration of AOPs with molecular datasets and slows progress towards quantitative AOP applications [1, 19]. Attaching a fitting GO BP annotation to each KE resolves this. Once a KE has an associated gene set, transcriptomics data can be mapped onto it and its perturbation scored, as Martens et al. showed using gene sets derived from WikiPathways [6]. A well-fitting term also narrows the analysis to the genes relevant to that KE, so that mechanistic signals are not diluted by the rest of the transcriptome.

Several studies have explored the use of omics data to quantify molecular events within AOP-related frameworks. Vuong et al. [20] integrated multi-omics data with benchmark dose modelling to establish dose-response relationships across KEs in acute myeloid leukaemia, demonstrating how transcriptomics links early molecular events to phenotypic outcomes. Martens et al. [6] developed molecular AOPs by mapping KEs to curated biological pathways in WikiPathways, then mapped transcriptomics data onto those pathways. Transcriptomics-based enrichment scores were calculated to assess KE perturbation in liver steatosis. Perkins et al. [21] integrated transcriptomics with causal networks to identify molecular signatures and pathway perturbations supporting chemical hazard assessment in rat liver cancer. Liu et al. [22] used time-concordant gene expression analysis to identify events preceding Drug-Induced Liver Injury (DILI), enabling mechanistic hypothesis generation.

These approaches relied on study-specific pathway mappings, curated gene lists, or expert-driven assignments tailored to particular AOPs or disease contexts. As the AOP framework expands to encompass diverse toxicological endpoints, there is a growing need for a systematic annotation methodology that ensures consistency and reproducibility across studies. The current study addresses this need by developing a systematic workflow for ontology-driven gene set mapping to KEs, establishing a generalizable annotation methodology applicable across different organs and toxicity types. We hypothesised that systematic ontological annotation would increase the number of KEs that can be linked to a gene set, and thereby the number of KEs whose perturbation can be measured from transcriptomics data. We annotated KEs across the brain, liver, lung, and kidney AOP networks, and revised the annotations that were already present. Afterwards, we evaluated the impact of our mapping by showing an increased number of transcriptomics-linkable KEs through a pesticide-induced liver toxicity case study, demonstrating how transcriptomics data can support KE-level perturbation analysis. Pesticides were chosen for this case study because the diet is the main route of human exposure [23]. The liver is where these compounds are metabolised and where these compounds result in adverse outcomes [24].

## 2 Methods

### 2.1 Ontology mapping for unannotated KEs

To enable transcriptomics-based assessment of AOPs, we systematically annotated unannotated KEs in AOP-Wiki using a structured curation workflow (Figure 1). All annotations were made by a single curator (S.A.) and reviewed by a second (M.M.) over several rounds, in which contested assignments were revised until agreement was reached on the best-fitting term.

**Fig. 1:**
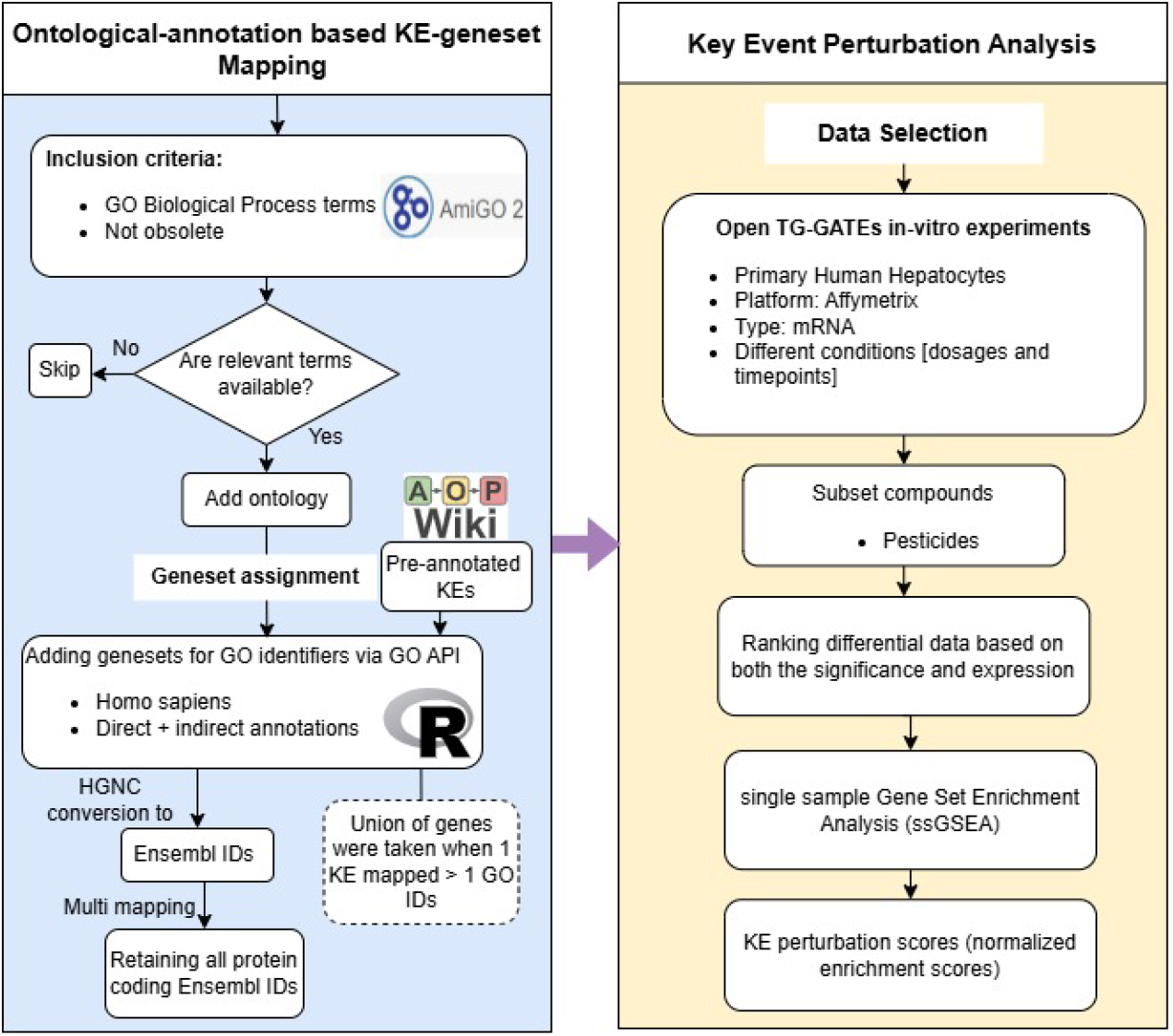
Workflow showing the decision tree to add GO BP terms to the KEs using information from KE-title and description (if available). Gene sets were added to the KEs based on the mapped GO identifier. These KEs with gene sets were then linked to Open TG-GATEs transcriptomics data of pesticide exposure across various conditions to analyse the KE perturbation patterns using single-sample gene set Enrichment Analysis (ssGSEA).

We focused on KEs without biological process annotations related to four target organs: brain, lung, liver, and kidneys. KEs were retrieved from the AOP-Wiki RDF SPARQL endpoint https://aopwiki.rdf.bigcat-bioinformatics.org/sparql/ using custom SPARQL queries (Supplementary File S1) based on the AOP-Wiki release accessed on 28 January 2025. Organ-specific AOPs were identified by querying AOP titles and adverse outcomes containing organ-specific keywords (given in respective SPARQL queries in the Supplementary File S1). Retrieved AOPs were subsequently manually filtered to exclude non-mammalian and multi-organ AOPs before extracting their associated KEs. For each unannotated KE, suitable Gene Ontology (GO) terms for biological processes were identified by extracting information from the KE title, description (if available), and relevant AOP context, using AmiGO2 https://amigo.geneontology.org/amigo as the search tool. To ensure comprehensive KE characterisation, biological object (BO) annotations were also added where appropriate (e.g., GO cellular component term ‘mitochondrion’ for KE 177 ‘Mitochondrial dysfunction’). During ontology mapping, directional terms (increase, decrease, activation, inhibition) were excluded to focus on the underlying biological process. The following filters were applied in AmiGO2: idspace = GO, is_obsolete = false, source = biological_process. Only *Homo sapiens*-specific gene identifiers were considered to ensure human-relevant annotations.

### 2.2 Improvements to the pre-annotated GO terms

Two curation steps were applied across all four organ networks. First, obsolete GO identifiers were replaced by their current valid equivalents so that the annotations remain resolvable against the live ontology. Second, BP terms that included directionality were replaced by direction-neutral terms; directionality (increase, decrease, agonism, inhibition) is recorded in the AOP-Wiki Action field and was therefore deliberately excluded from the GO term. Both sets of replacements are listed in Supplementary Table 2.

For the liver network, the BP annotations of the pre-annotated KEs were additionally reviewed manually against the current release of the Gene Ontology. The annotations retrieved from AOP-Wiki were frequently uninformative: many KEs carried high-level parent classes (for example *transport, metabolic process, signaling, biosynthetic process*) that resolve to large, non-specific gene sets. To address this, each existing annotation was reviewed and, where a more relevant GO BP class was available, replaced by the better-fitting term. Annotations that already described the event well were retained unchanged. Terms were anchored on the biological entity named in the KE, so that entity-specific classes were preferred over generic ones (e.g. KE 41, Inhibition, Bile Salt Export Pump (ABCB11), was assigned *canalicular bile acid transport* in place of the generic *transport*). Where a KE bundled two distinct processes, up to two co-equal terms were assigned and their gene sets pooled as a union. A small number of KEs carry Molecular Function rather than Biological Process terms; these were left unchanged. In AOP-Wiki events at the molecular level (MIEs) are described by molecular functions. All revisions are listed in Supplementary Table 3.

### 2.3 GO-based gene set assignment to KEs

Gene sets were assigned to both pre-annotated and newly-annotated KEs based on their GO identifiers. An R script (version 4.4.2, session information is available in the GitHub repository) retrieved human gene annotations via the GO API, including both direct and inferred (parent–child) annotations to capture comprehensive gene-process relationships. The annotations obtained from the GO API were provided in HGNC symbol format. To enable integration with other datasets and ensure standardised identifiers, these HGNC symbols were converted to Ensembl gene identifiers using the `BridgeDbR` R package (2.16.0) with the *Homo sapiens* mapping file `Hs_Derby_Ensembl_111.bridge` (Ensembl release 111). When a single HGNC symbol mapped to multiple Ensembl IDs, all protein-coding IDs were retained to ensure comprehensive coverage and account for assembly updates, duplicated loci, or alternative haplotype sequences.

### 2.4 AOP network construction and data visualisation

Each organ-specific AOP network consists of KEs as nodes and KERs as edges. We used the KEs retrieved in Section 2.1, and obtained their KERs with further SPARQL queries (Supplementary File S1). Networks were visualised in Cytoscape (v3.10.3) using the RCy3 R package (v2.26.0).

KE–gene set mappings were added as node attributes. Nodes were coloured by annotation status (pre-annotated / newly-annotated) and sized in proportion to the number of genes in the associated gene set.

We then tested the effect of this annotation in a case study. The liver network was used to score KE perturbation from transcriptomics data of pesticide-exposed human hepatocytes, as described in the following sections.

### 2.5 Liver case study: AOP network

The liver was chosen for the case study because it is the primary site of xenobiotic metabolism and therefore the organ most exposed to ingested pesticides, and because human hepatocyte transcriptomics data covering a range of compounds are publicly available through Open TG-GATEs. The liver AOP network was constructed as described in Section 2.4. Its pre-annotated KEs were additionally revised (Section 2.2), so this network carries a third annotation status that the other organ networks do not. Nodes were therefore coloured by four categories: pre-annotated, improved pre-annotated, newly-annotated, and KEs without an associated gene set.

#### 2.5.1 KE grouping by GO annotation and directionality

A GO BP term is often the most fitting class available for more than one KE, so KEs that describe different events can carry the same annotation. For example, KE 89 ‘Synthesis, De Novo Fatty Acid (FA)’, KE 458 ‘Increased, De Novo FA synthesis’, and KE 463 ‘Up Regulation, FAS’ are all annotated with ‘fatty acid biosynthetic process’ (GO:0006633). This reflects the granularity of the Gene Ontology rather than redundancy among the KEs. Such KEs share a gene set and therefore receive identical perturbation scores, so the transcriptomics readout cannot separate them. They were grouped for this reason, and the resulting groups are used when reporting perturbed KEs (Section 2.7.3).

To group similar KEs in the liver AOP network, KEs were grouped by their GO annotation together with the direction of change. The direction was derived from the action term of each KE. Action terms were retrieved from the AOP-Wiki RDF (predicate pato:0000001) through its SPARQL endpoint. KEs that carried no action term in the AOP-Wiki were assigned one in the present study on the basis of their title, using the same vocabulary as the AOP-Wiki, and the origin of every action term was recorded in a separate column (AOP-Wiki RDF or current study). The eleven action terms in use were then translated into four directions: ‘+’ for an increase (increased, premature), ‘-’ for a decrease (decreased, arrested, delayed), ‘Change’ for a qualitative change without a sign (abnormal, disrupted, functional change, morphological change, pathological), and ‘Occurrence’ for events that simply take place (occurrence). ‘Change’ was kept separate from ‘Occurrence’ because these terms do state that something changed, only not in which direction. A KE with several KECs carries several action terms; when only one of them was signed, that term determined the direction, and the single KE whose terms were in conflict was resolved manually. KEs were finally grouped when they shared the same direction and exactly the same set of GO identifiers; for KEs annotated with more than one GO term, all identifiers had to match. Grouping in this way consolidates KEs that describe the same biological process changing in the same way.

### 2.6 Transcriptomics data selection and processing

To assess the revised liver AOP network, publicly available transcriptomics data were selected for KE perturbation analysis. The Open TG-GATEs human dataset was retrieved using the ToxicoGx R package (v2.10.0), containing gene expression data from primary human hepatocytes (PHH) exposed to 146 chemical compounds. To focus on regulatory-relevant substances, we filtered for pesticides, yielding four compounds: cycloheximide (fungicide [25]), 2,4-dinitrophenol (herbicide/insecticide [26]), hexachlorobenzene (fungicide [27]), and rotenone (insecticide [28]). The expression data provided by Open TG-GATEs are already log-normalised. Each compound was tested at three concentrations (low, middle, high) in two biological replicates, with time-matched vehicle controls. Cycloheximide and 2,4-dinitrophenol were tested at 4, 20 and 100 µM, rotenone at 0.08, 0.4, and 2 µM, and hexachlorobenzene at 1.2, 6 and 30 µM. Exposures lasted 8 and 24 h for all compounds, with an additional 2 h time point for hexachlorobenzene. This gave six experimental conditions per compound and nine for hexachlorobenzene. Differential expression analysis was performed for each compound and condition using the `limma` R package (v3.62.2), comparing treated samples relative to controls. For each experimental condition, log_2_FC values and p-values were exported for downstream KE perturbation analysis.

### 2.7 KE perturbation analysis

#### 2.7.1 Handling multiple GO Annotations per KE

Some pre-annotated KEs in AOP-Wiki were linked to multiple GO terms due to complex biological processes involving several mechanisms (e.g., KE 853: ‘Changes/Inhibition, Cellular Homeostasis and Apoptosis’ annotated with GO:0019725 and GO:0006915) or differing author perspectives (e.g., KE 768: ‘Increase, Cytotoxicity’ annotated with GO:0008219 and GO:0001909). These GO terms may be related through *is_a* or *part_of* relationships. To preserve biological complexity while enabling quantification, we took the union of gene sets from all associated GO terms for each KE.

#### 2.7.2 Computing KE perturbation scores

KE perturbation scores were computed using single-sample gene set Enrichment Analysis (ssGSEA) [29]. Gene sets containing 10–800 genes were included in the analysis. The lower threshold was applied to exclude very small gene sets that may yield unstable enrichment estimates. The upper threshold was increased beyond commonly used default thresholds to retain biologically relevant GO-based KE gene sets that are inherently larger. For each KE, genes were ranked by a combined score: log_2_FC × (–log_10_(*p*-value)), integrating both effect size and statistical confidence. A running sum (RS) was computed by walking down the ranked list, incrementing RS when encountering KE gene set genes (weighted by |log_2_FC| ^*α*^, where *α* = 0.25) and decrementing for non-gene set genes. *α* is the weighting parameter that controls the contribution of the absolute log_2_ fold changes (|log_2_ FC|) to the enrichment score. When *α* = 0, all genes in the gene set contribute equally (unweighted enrichment). We set *α* = 0.25 to give slightly greater weight to genes with larger absolute fold changes while preventing genes with extremely large fold changes from dominating the enrichment score.

The enrichment score (ES) was defined as the maximum absolute deviation of RS from zero. To account for gene set size and expression distribution, the observed ES was normalised against a null distribution generated through 1,000 random permutations of equally sized gene sets, yielding a Normalised Enrichment Score (NES). Statistical significance was assessed via permutation testing with one-tailed *p*-values, calculated as the proportion of permuted scores equal to or more extreme than the observed score. A random seed (123) ensured reproducibility. KEs with |NES| *>* 1.5 and *p <* 0.05 were classified as significantly perturbed.

#### 2.7.3 Comparison of perturbation profiles across compounds

The liver network was assessed in State III, which combines the revised pre-annotated KEs with the newly-annotated ones. KEs belonging to the same group (Section 2.5.1) were first collapsed into a single row, since group members carry the same GO terms and therefore the same gene set and the same NES.

##### Principal component analysis

NES values were arranged in a matrix with one row per condition (a compound at a given concentration and time point) and one column per KE or KE group. Missing values were set to zero, and KEs with the same value in every condition were removed as uninformative. The matrix was centred and scaled, and principal component analysis was performed with `prcomp` (`R` stats package). The first two principal components were plotted with one point per condition, coloured by compound and shaped by dose, with an ellipse covering 80% of the conditions of each compound. Hexachlorobenzene was measured at three time points and the other three compounds at two, giving 27 conditions in total; hexachlorobenzene therefore contributes more conditions than the other compounds, and its ellipse is expected to cover a correspondingly wider area. To judge whether the compounds separate more than their own conditions vary, the mean distance of each condition to the centre of its own compound and the distances between the compound centres were calculated in the PC1–PC2 plane.

##### UpSet plot

A KE or KE group was counted as perturbed for a compound when it met the significance criteria in either direction in at least one of that compound’s conditions. This gave a binary matrix of KEs by compounds, which was visualised as an UpSet plot with the UpSetR R package (v1.4.0), showing the largest intersections ordered by size. The number of compounds in which each KE was perturbed was tabulated to identify the KEs perturbed by all four compounds and those perturbed by a single compound.

## 3 Results

### 3.1 AOP network generation

The AOP networks for brain, kidney, liver, and lung comprise 203, 68, 196, and 134 KEs, respectively (see Table 1). Detailed information on the top 5 hub KEs, including degree and betweenness centrality, is provided in Supplementary Table 1. The majority of KEs are organ-specific, with brain having the highest number of unique KEs (169), followed by liver (163), lung (111), and kidney (45) (Figure 2). Only three KEs are shared among all four networks.

**Table 1:** Organ-specific AOP networks showing the number of associated AOPs, total KEs, and hub KEs. Six KEs are listed for liver because KE 57 and KE 142 share the same degree and betweenness centrality and therefore rank jointly fifth.

| Organ | AOPs | Total distinct KEs | MIEs | AOs | Top 5 hub KEs |
| --- | --- | --- | --- | --- | --- |
| Brain | 47 | 203 | 43 | 24 | KE 1393, KE 341, KE 188, KE 352, KE 55 |
| Kidney | 16 | 68 | 15 | 7 | KE 177, KE 814, KE 1097, KE 1392, KE 1115 |
| Liver | 36 | 196 | 45 | 18 | KE 177, KE 459, KE 373, KE 1289, KE 57, KE 142 |
| Lung | 27 | 134 | 20 | 17 | KE 1496, KE 1457, KE 1115, KE 1392, KE 68 |

**Fig. 2:**
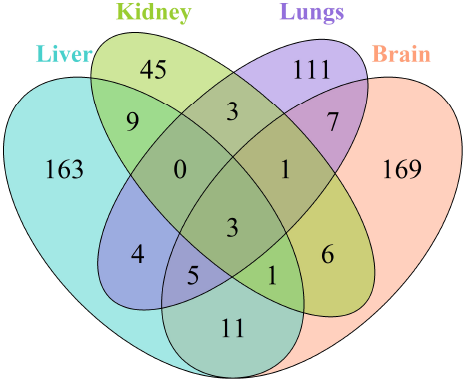
Overlap of KEs across the liver, kidney, lung, and brain AOP networks. The limited overlap indicates that most KEs are organ-specific, with only three KEs shared among all four networks.

### 3.2 GO identification

Analysis of organ-specific KEs from AOP-Wiki revealed that 56.7% of those in the brain, kidney, liver and lung networks already carried a biological process (BP) annotation (Figure 3). Notably, KEs at lower biological organisation levels (molecular, cellular, and tissue) are predominantly annotated with Gene Ontology (GO) BP terms. We systematically mapped GO BP terms to unannotated KEs (shown as white bars in Figure 3) using the curation workflow described in the Methods section (Figure 1). The resulting annotations for the brain, kidney, liver, and lung AOP networks have been made publicly available through Zenodo [30].

**Fig. 3:**
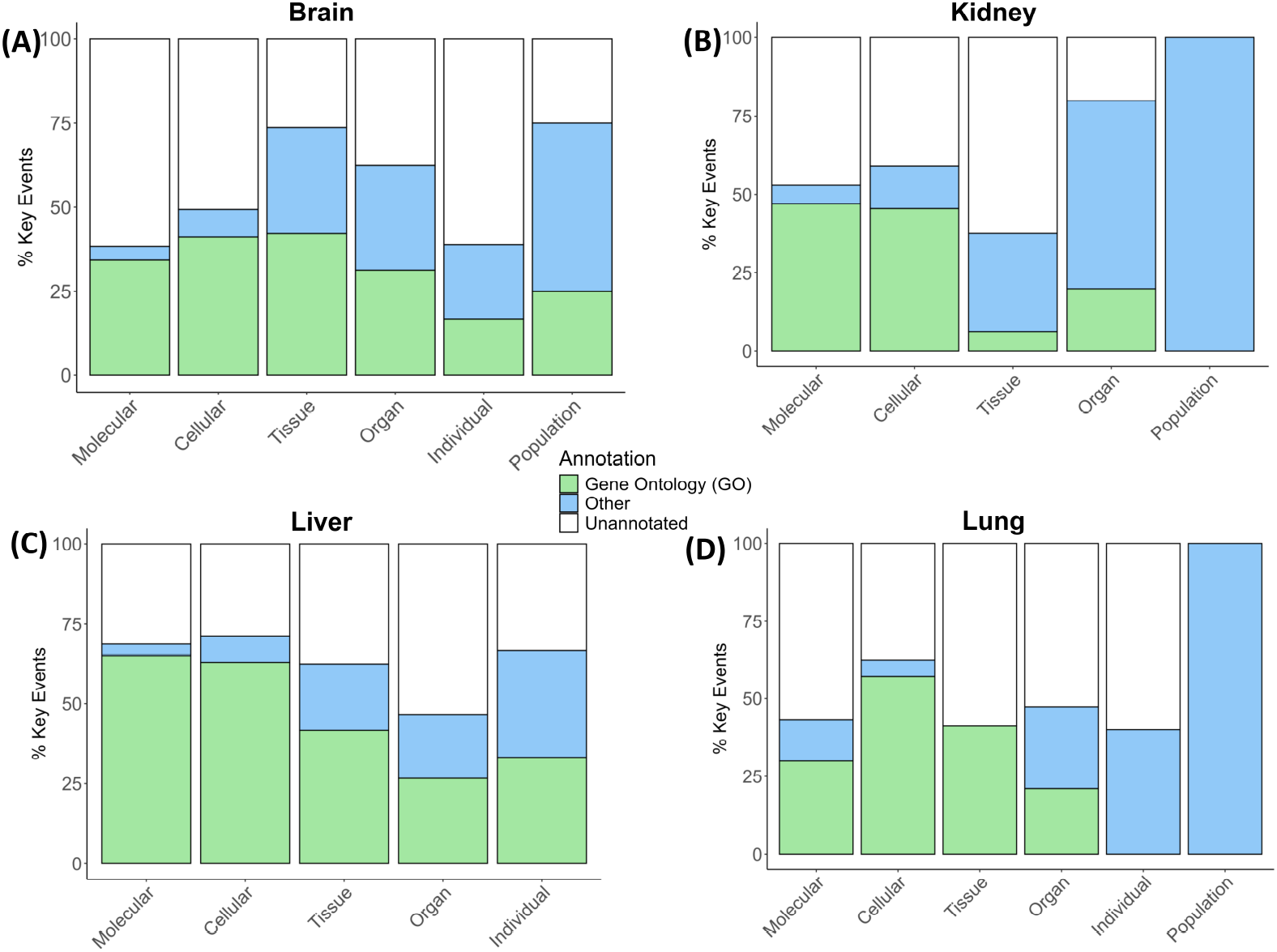
Annotation status of Key Events across biological organisation levels in four organ systems. Stacked bar plots showing the percentage of Key Events (KEs) annotated with Gene Ontology (GO) terms (green), other ontologies (blue), or remaining unannotated (white) at each biological organisation level (Molecular, Cellular, Tissue, Organ, Individual, Population) for (A) Brain, (B) Kidney, (C) Liver, and (D) Lung.

### 3.3 Ontology mapping for unannotated KEs

A GO BP annotation is what allows a gene set to be attached to a KE, and therefore what makes that KE linkable with transcriptomics. To evaluate how much our ontology-driven annotation improved this, we compared State I (pre-annotated KEs only) with State II (pre-annotated & newly-annotated KEs) across the four organ networks (Figure 4). GO term coverage analysis (Figure 4(C) & (D)) demonstrated organ-specific expansion of molecular annotations. Median gene set sizes decreased in State II relative to State I for brain, liver and lung (Table 2). Liver showed the largest reduction (-67.3%), followed by lung (-65.3%) and brain (-51.7%). Kidney was the exception, with a median gene set size that was essentially unchanged (+1 gene, +1.3%). The reduction in the other three organs arises because the newly-annotated KEs carry more fitting, and therefore smaller, GO terms than the pre-existing annotations. Kidney had the smallest State I median of the four organs (75 genes), so its pre-existing annotations were already reasonably fitting. Overall, the percentage of KEs with associated gene sets increased after our curation effort in State II (Table 3).

**Fig. 4:**
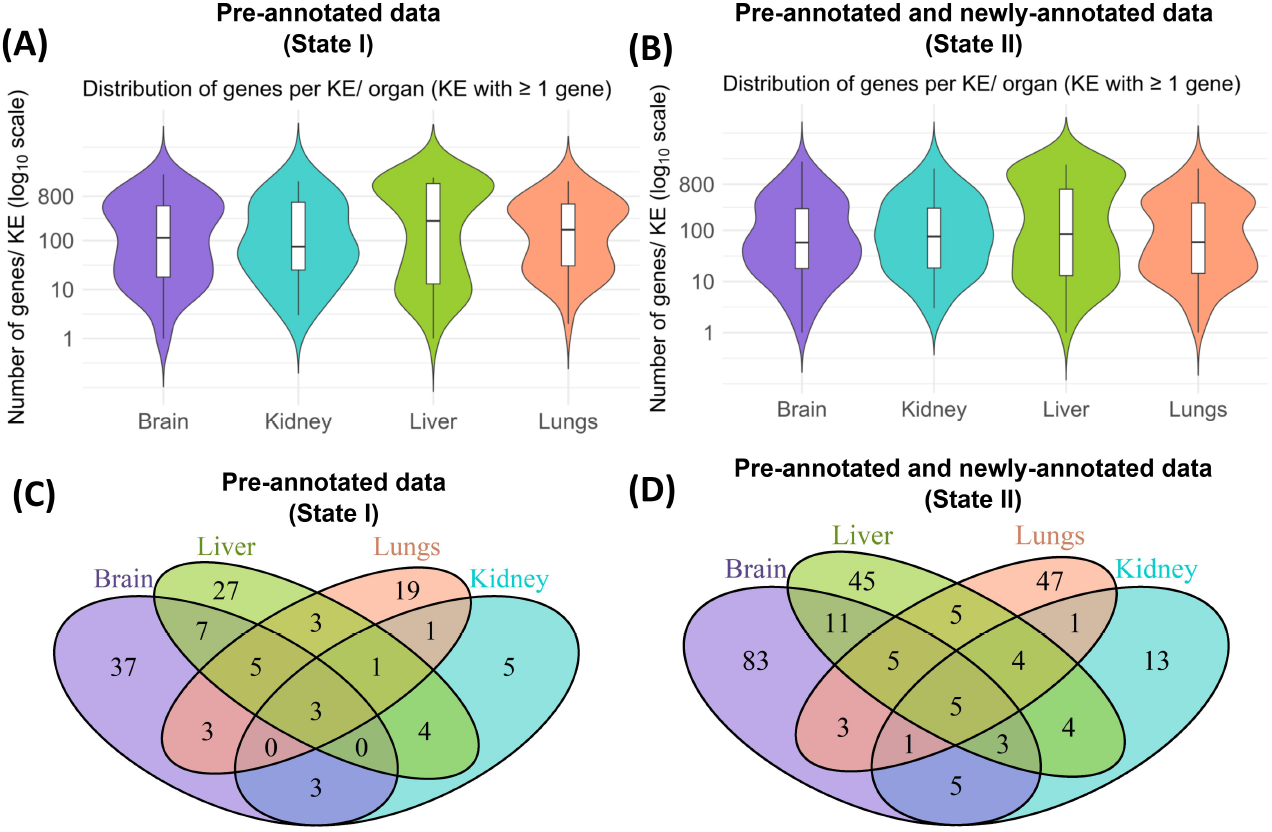
(A) Distribution of genes per KE across organs in pre-annotated (State I) AOP-Wiki data. (B) Distribution of genes per KE across organs after ontological annotation-based mapping (pre-annotated and newly-annotated data combined (State II)). (C) Overlap of GO terms across organ-specific networks in pre-annotated data (State I). (D) Overlap of GO terms across organ-specific networks after ontological annotation (pre-annotated and newly-annotated data combined (State II)).

**Table 2:**
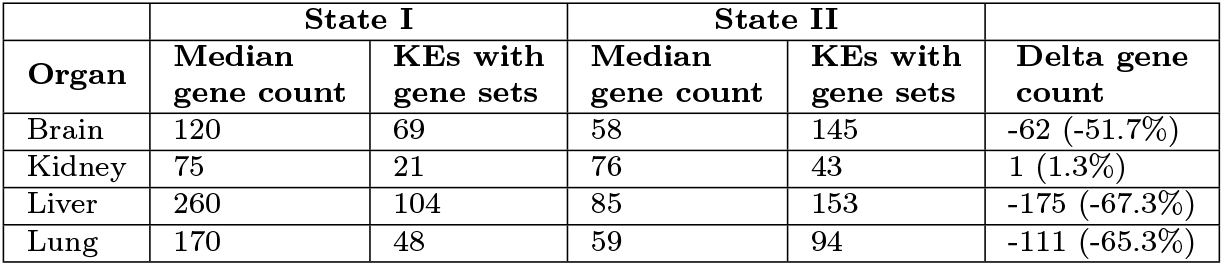
Comparison of median gene set sizes between State I (pre-annotated) and State II (pre-annotated + newly-annotated). Delta represents the absolute change in median gene set size between State I and State II, while delta % expresses this change relative to State I.

| Organ | State I |  | State II |  | Delta gene count |
| --- | --- | --- | --- | --- | --- |
|  | Median gene count | KEs with gene sets | Median gene count | KEs with gene sets |  |
| Brain | 120 | 69 | 58 | 145 | -62 (-51.7%) |
| Kidney | 75 | 21 | 76 | 43 | 1 (1.3%) |
| Liver | 260 | 104 | 85 | 153 | -175 (-67.3%) |
| Lung | 170 | 48 | 59 | 94 | -111 (-65.3%) |

**Table 3:** KEs with gene sets in State I (pre-annotated) and State II (pre-annotated & newly annotated). For liver, State III (improved pre-annotated data along with newly-annotated data) is also shown.

| Organ | Total KEs | State I<br>KEs with gene sets | State II<br>KEs with gene sets | State III (liver only)<br>KEs with gene sets |
| --- | --- | --- | --- | --- |
| Brain | 203 | 69 (34%) | 145 (71%) | - |
| Kidney | 68 | 21 (31%) | 43 (63%) | - |
| Liver | 196 | 104 (53%) | 153 (78%) | 151 (77%) |
| Lung | 134 | 48 (36%) | 94 (70%) | - |

### 3.4 Impact of revising the pre-annotated GO terms

Across the four organ networks, nine obsolete GO terms were replaced by their current equivalents, and three KEs (KE 115, KE 1015 and KE 2242) carried terms that included directionality and were replaced by direction-neutral terms (Supplementary Table 2).

For the liver network, 105 pre-annotated KEs were assessed. Fifty-five retained their original annotation, and 50 were revised (Supplementary Table 3). One of these revisions is a removal rather than a replacement: KE 18 carried an obsolete term for which no suitable replacement could be identified, so it was left without a Biological Process annotation and has no gene set. Three of the revisions are label drift rather than re-annotation: the GO identifier is unchanged, but the term has since been relabelled in the Gene Ontology while the AOP-Wiki record retains the superseded label. These are KE 57 and KE 854, for which GO:0008283 has been relabelled from *cell proliferation* to *cell population proliferation*, and KE 202, for which GO:0007249 has been relabelled from *I-kappaB kinase/NF-kappaB signaling* to *canonical NF-kappaB signal transduction*. The revised terms return smaller and more specific gene sets. KE 454 (*Increased, Triglyceride formation*), for example, changed from *biosynthetic process* (GO:0009058, 1,738 genes) to *triglyceride biosynthetic process* (GO:0019432, 19 genes). Gene sets were assigned to 104 of the 196 liver KEs (53%) in State I, 153 (78%) in State II and 151 (77%) in State III (Table 3). The decrease from State II to State III is accounted for by two KEs, KE 167 and KE 483, which were assigned terms to which no human genes are currently annotated and therefore have no gene set in State III. The median gene set size in the liver network fell from 260 genes in State I to 40 in State III (n = 151), a reduction of 220 genes (84.6%).

### 3.5 Case study

#### 3.5.1 AOP liver network

The effect of the annotations becomes visible when they are mapped back onto the network they came from. Figure 5 shows the liver AOP network in State III, in which 151 of the 196 KEs (77%) carry a gene set. The 45 grey nodes are the KEs that still lack a Biological Process annotation and therefore remain inaccessible to transcriptomics; these sit predominantly at the organ, individual, and population levels, where the events described are not gene expression processes. Overall, the network provides a more relevant basis for transcriptomics-based assessment of KE perturbation.

**Fig. 5:**
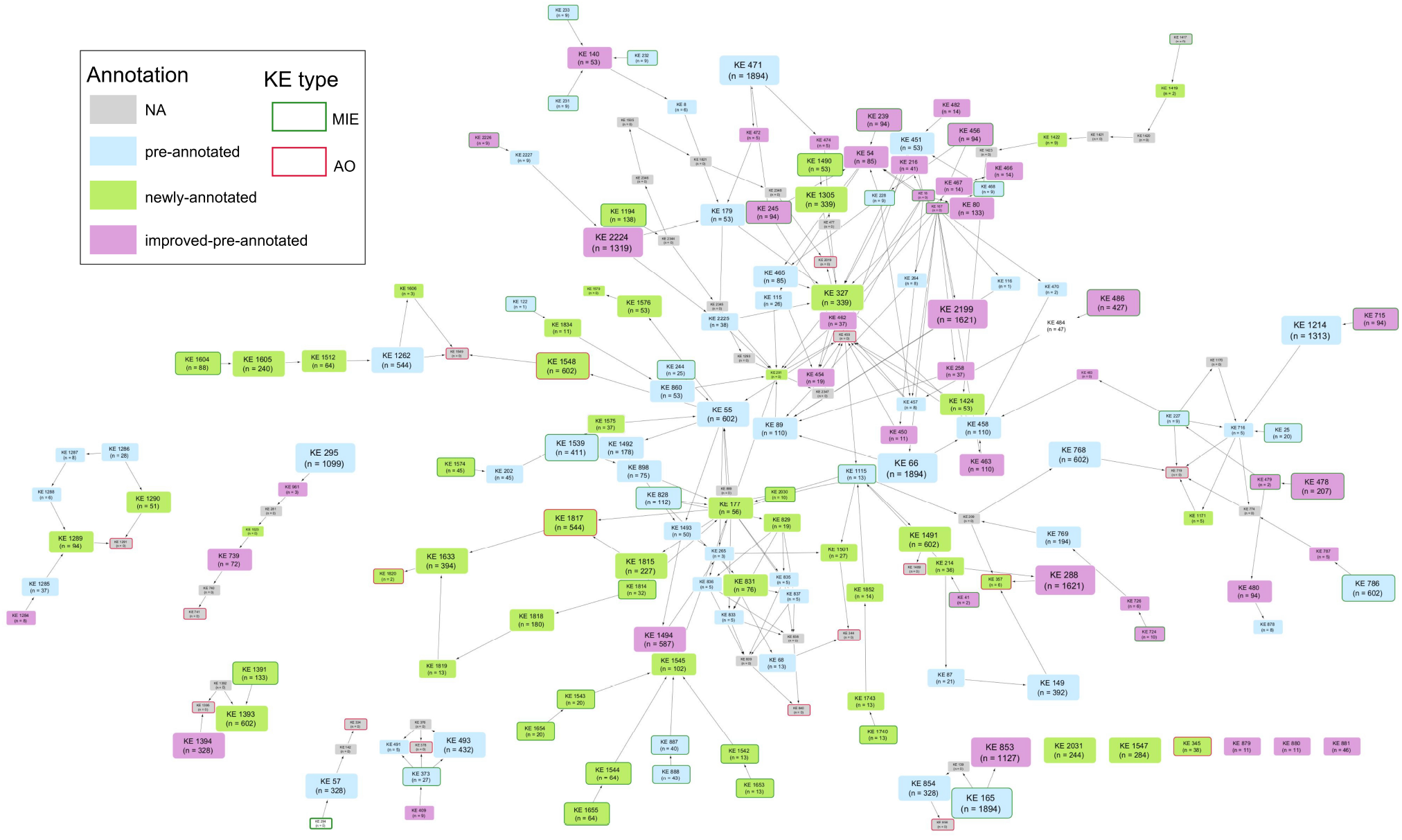
The liver AOP network in State III (improved-pre-annotated combined with newly-annotated). Nodes represent key events (KEs) and directed edges represent key event relationships (KERs). Node colour indicates annotation status: blue for KEs pre-annotated with a Gene Ontology (GO) Biological Process (BP) term in AOP-Wiki, purple for KEs whose pre-existing annotation was improved in this study, green for KEs newly-annotated in this study, and grey for KEs that remain without a GO BP annotation. Node size is proportional to the number of genes in the associated gene set, with grey nodes drawn at a fixed minimum size. In total, 151 of the 196 KEs (77%) carry a gene set in this state.

##### 3.5.2 KE perturbation patterns across pesticides

Of the 151 KEs carrying a gene set in State III, 101 had a gene set of between 10 and 800 genes and were taken forward for scoring; 41 were excluded as too small (fewer than 10 genes) and 9 as too large (more than 800 genes). Applying the KE grouping to these 101 KEs gave 73 KE units for the analysis: 17 groups covering 45 KEs, and 56 KEs that belong to no group. Of these 73 KE units, 67 were perturbed by at least one compound. The remaining 6 KE units, covering 7 KEs, were not perturbed by any compound. The number of perturbed KE units per compound was 57 for cycloheximide, 52 for hexachlorobenzene, 51 for 2,4-dinitrophenol and 29 for rotenone.

In the principal component analysis of the individual conditions, the first two components explained 32.5% and 17.5% of the variance (50.0% together) (see Figure 6(A)). Cycloheximide separated from the other three compounds along PC1, while the conditions of 2,4-dinitrophenol, hexachlorobenzene and rotenone overlapped around the origin. The mean distance of a condition to the centre of its own compound was 5.61 for hexachlorobenzene, 3.21 for cycloheximide, 3.18 for 2,4-dinitrophenol and 1.93 for rotenone. Hexachlorobenzene contributed nine conditions against six for the other compounds, which accounts for part of its wider spread. The distances between the compound centres were 8.11 to 10.55 for cycloheximide against each of the other compounds, and 1.14 to 2.82 among 2,4-dinitrophenol, hexachlorobenzene and rotenone. Only cycloheximide was therefore separated by more than the spread of the conditions within a compound.

**Fig. 6:**
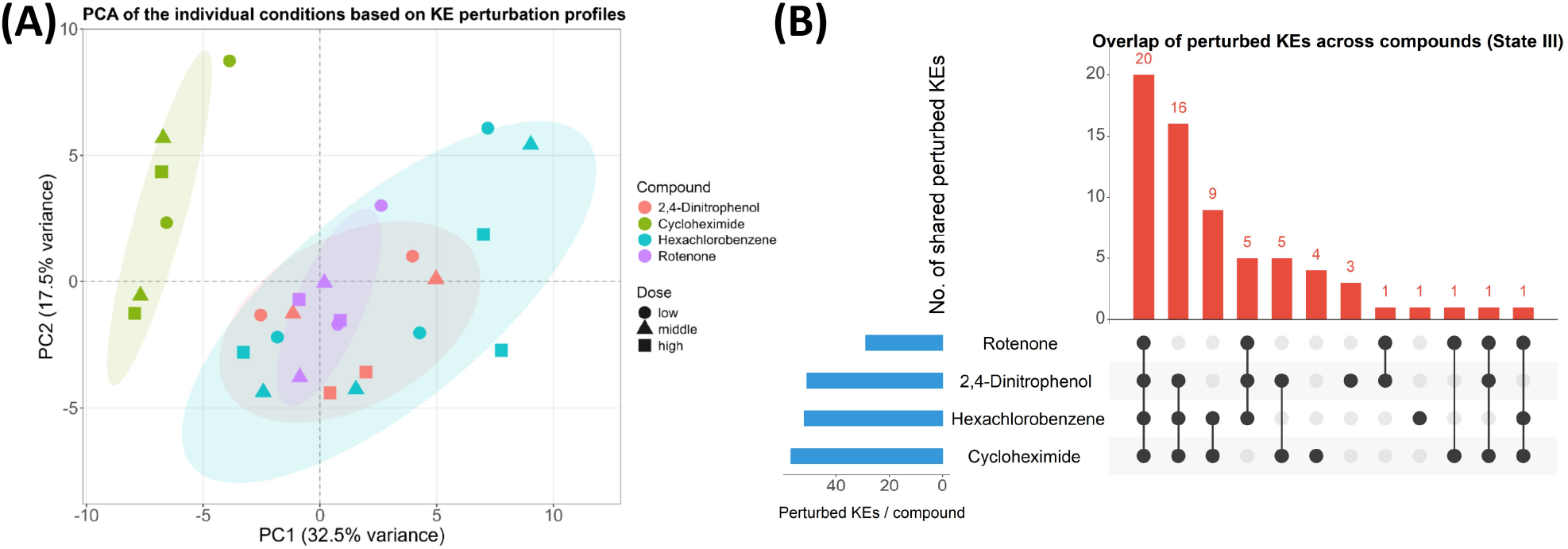
(A) Principal component analysis (PCA) of the 27 individual exposure conditions, based on the normalised enrichment score (NES) of 73 KE units (56 single KEs and 17 KE groups). Each point is one condition (compound at a given dose and time point), coloured by compound and shaped by dose; ellipses cover 80% of the conditions of a compound. PC1 and PC2 together explain 50.0% of the variance. Hexachlorobenzene was measured at three time points (2, 8 and 24 hours) and the other compounds at two (8 and 24 hours), so it contributes nine conditions against six for the others and its ellipse covers a wider area. (B) UpSet plot of the KEs perturbed by each compound. A KE was counted as perturbed when it met the significance criteria in at least one condition of that compound. The upper bars give the number of KEs in each intersection, the left bars the number of perturbed KEs per compound.

The two components were driven by different sets of processes. PC1 was loaded mainly by KEs of lipid and xenobiotic metabolism, including protein N-linked glycosylation (KE 1604), triglyceride biosynthetic process (KE 454), bile acid and bile salt transport (KE 214), very-low-density lipoprotein particle assembly (KE 879; KE 880) and fatty acid beta-oxidation (KE 1490; KE 1424; KE 860; KE 140; KE 179; KE 451). PC2 was loaded mainly by KEs of mitochondrial organisation and DNA damage, including organelle disassembly (KE 898), protein alkylation (KE 244), mitochondrial membrane organisation (KE 177), DNA repair (KE 493) and signal transduction in response to DNA damage (KE 1194).

While the principal component analysis describes how the conditions relate to one another, the perturbed KEs themselves are compared across compounds in Figure 6(B). Twenty KE units were perturbed by all four compounds, 23 by three, 16 by two and 8 by a single compound. The largest intersection was the set shared by all four compounds. Compound-specific perturbations were few: 4 KE units for cycloheximide, 3 for 2,4-dinitrophenol, 1 for hexachlorobenzene and none for rotenone.

These 20 KE units, which together represent 24 KEs, are listed in Table 4. They fall into five functional categories: mitochondrial function and energy metabolism accounts for nine KEs, lipid metabolism and transport for six, and DNA damage response and cell death, tissue remodelling and fibrosis, and inflammation and immune signalling for three each. All 20 could be assigned to a category. Two biological processes appeared twice in the table, as separate units with opposite directions. Very-low-density lipoprotein particle assembly (GO:0034379) is carried by KE 879 and KE 880, grouped with direction ‘+’, and by KE 450, with direction ‘-’. Angiotensin maturation (GO:0002003) is carried by KE 1743 with direction ‘+’ and by KE 1740 with direction ‘Change’. In both cases, the KEs share the GO term but differ in direction and were therefore not grouped.

**Table 4:** KEs consistently perturbed across all four compound exposures. Of the 73 KE units scored in the liver network (56 single KEs and 17 KE groups), 20 were perturbed by every compound. These were assigned to five functional categories based on their associated biological process terms to facilitate biological interpretation: mitochondrial function and energy metabolism, lipid metabolism and transport, DNA damage response and cell death, tissue remodelling and fibrosis, and inflammation and immune signalling. All 20 could be assigned to a functional category. KE IDs separated by a semicolon denote KEs that were grouped because they share the same GO biological process term and direction.

| Sr. no. | KE ID | GO term | Functional group |
| --- | --- | --- | --- |
| 1 | KE 493 | DNA repair | DNA damage response and cell death |
| 2 | KE 2031 | apoptotic signaling pathway | DNA damage response and cell death |
| 3 | KE 1194 | signal transduction in response to DNA damage | DNA damage response and cell death |
| 4 | KE 149;<br>KE 1633 | inflammatory response | Inflammation and immune signalling |
| 5 | KE 1819 | type II interferon-mediated signaling pathway | Inflammation and immune signalling |
| 6 | KE 1289 | cholesterol homeostasis | Lipid metabolism and transport |
| 7 | KE 54;<br>KE 465 | fatty acid transport | Lipid metabolism and transport |
| 8 | KE 879;<br>KE 880 | very-low-density lipoprotein particle assembly | Lipid metabolism and transport |
| 9 | KE 450 | very-low-density lipoprotein particle assembly | Lipid metabolism and transport |
| 10 | KE 887 | NADH dehydrogenase (ubiquinone) activity | Mitochondrial function and energy metabolism |
| 11 | KE 769 | cellular respiration | Mitochondrial function and energy metabolism |
| 12 | KE 888 | mitochondrial electron transport, NADH to ubiquinone | Mitochondrial function and energy metabolism |
| 13 | KE 177 | mitochondrial membrane organisation | Mitochondrial function and energy metabolism |
| 14 | KE 1547 | mitochondrion organisation | Mitochondrial function and energy metabolism |
| 15 | KE 898 | organelle disassembly | Mitochondrial function and energy metabolism |
| 16 | KE 1545 | oxidative phosphorylation | Mitochondrial function and energy metabolism |
| 17 | KE 1544;<br>KE 1655 | proton motive force-driven mitochondrial ATP synthesis | Mitochondrial function and energy metabolism |
| 18 | KE 1743 | angiotensin maturation | Tissue remodelling and fibrosis |
| 19 | KE 1740 | angiotensin maturation | Tissue remodelling and fibrosis |
| 20 | KE 1501 | extracellular matrix assembly | Tissue remodelling and fibrosis |

The 67 perturbed KE units comprised 94 KEs. The additional annotation determined what could be observed among them. Of the 94 KEs perturbed in State III, 39 were newly-annotated and 26 were improved, so most of the response would have been invisible or differently defined in the pre-annotated network. The same holds for the KEs perturbed by all four compounds: only 7 of 24 were pre-annotated.

## 4 Discussion

We aimed to improve gene-to-KE mapping through ontology annotation, enabling more KEs to be assessed using gene-expression data. When the BP component of a KE is described by a GO BP term, it provides the basis for mapping a gene set to that KE. However, at present, more than half (52%) of the KEs in the AOP-Wiki are not described by BP annotations. This incomplete annotation hinders the mapping of transcriptomics data onto those KEs and the assessment of KE perturbation. To address this gap, we systematically mapped GO BP terms to previously unannotated KEs in four organs, thereby increasing the proportion of KEs that can be linked to transcriptomics data and assessed for perturbation.

The AOP network generation revealed that the selected organ networks capture largely distinct mechanistic pathways leading to tissue-specific adverse outcomes, as the majority of KEs were organ-specific with minimal overlap across organs. Only ‘Mitochondrial dysfunction’ (KE 177), ‘Reactive oxygen species’ (KE 1115), and ‘Oxidative stress’ (KE 1392) were shared among all four organs (Figure 2), consistent with their well-established role in toxicity [31, 32].

Analysis revealed that 57% of KEs related to four organ systems were already annotated with BP terms, while the remaining 43% required new annotations. Notably, KEs at lower biological organisation levels (molecular, cellular, and tissue) showed higher annotation rates with GO terms (Figure 3). The reason is structural. GO describes what gene products do and contains no terms for events at the level of an organ, an individual or a population. This distribution is advantageous for transcriptomics integration, because molecular and cellular KEs represent the mechanistic events most directly measurable via changes in gene expression.

We observed systematic changes in gene set size distribution across all organs after applying our ontology mapping strategy (Figure 4). In State I, broad density distributions reflected heterogeneous gene-to-KE mappings characteristic of existing AOP-Wiki annotations (Figure 4(A)). Following our systematic curation in State II, density peaks shifted toward lower gene counts in brain, liver and lung, because the newly-annotated KEs were assigned more fitting biological process terms, which in most cases were more specific than those already present in AOP-Wiki (Figure 4(B)). In kidney, the median was unchanged, although the upper part of the distribution still came down (Table 2). Kidney is the smallest of the four networks and its pre-existing annotations were already fitting, so there was little scope for further reduction in gene set size. Notably, the most relevant term is not always the most specific, as some KEs require broader terms than others. Therefore, we mapped each KE to the most appropriate GO BP term(s), selecting either specific (lower-level) or broad (higher-level) terms based on the semantic fit and biological granularity of the KE.

GO term coverage analysis revealed that the number of unique GO terms rose from 118 to 235 across the four networks. Brain gained most in absolute terms (58 to 116, +58), followed by lung (35 to 71, +36), liver (50 to 82, +32) and kidney (17 to 36, +19) (Figure 4(C) & (D)). The proportion of GO terms shared between organs fell from 25% to 20%, consistent with the organ-specific nature of the toxicological pathways (Figure 4 (C) & (D)). As a whole, the percentage of KEs with gene sets, and therefore linkable to transcriptomics, increased after ontology-driven KE-to-gene set mapping within each organ system (Table 3).

Together, these results suggest that systematic ontology-driven annotation improves coverage and precision in different ways. The breadth of molecular coverage across organ-specific AOP networks (more unique GO terms per organ) increased in all four organs, as did the number of KEs carrying a gene set, because both depend on how many KEs lacked an annotation rather than on the quality of those already present. The precision of individual KE-gene associations (smaller, more specific gene sets) improved only where the pre-existing annotations were broad: the median fell from 260 to 85 genes in liver, 170 to 59 in lung and 120 to 58 in brain, while kidney, which had the smallest gene sets to begin with, did not change (Table 2). Annotation narrows a gene set only when the term already in place is wider than the one that fits the KE.

Revising existing annotations proved as substantial a task as adding new ones: nearly half of the pre-annotated liver KEs required revision. Not all of these were corrections. In three cases, only the label had drifted while the identifier remained valid. AOP-Wiki stores the label alongside the identifier and does not update it when the ontology changes, so an annotation can look outdated while remaining correct. Storing the identifier and resolving the label at query time would remove this problem entirely, and would apply to every ontology AOP-Wiki draws on, not only GO. Revision also costs coverage. A GO term can describe a KE correctly and still return nothing measurable, because gene-to-term annotation coverage is itself incomplete. This is a limitation that curation effort cannot remove, and one that will lift as the Gene Ontology grows without any change to the AOP. Liver is the only network in which the pre-existing annotations were also revised, so this reduction combines the effect of adding new annotations with that of replacing overly general terms. The net effect for the liver network was a small loss of coverage in exchange for more specific gene sets. This results in a more relevant AOP network for biological interpretation, because each perturbation score now comes from a term that matches what the KE describes.

GO BP annotations themselves are not biomarkers for KEs, but the gene sets they link to serve as approximations of KE-associated molecular signatures. Multiple KEs can share the same GO BP term. For instance, the GO term ‘cell population proliferation’ (GO:0008283) is mapped to several KEs: KE 1394 (Induction, persistent proliferation/sustained proliferation), KE 57 (Pro-liferation, cell proliferation in the absence of cytotoxicity), and KE 739 (Increase, hypertrophy and proliferation (follicular cell)). The genes in this category therefore do not uniquely identify a specific KE, as GO BPs are functional categories rather than mechanism-specific markers. This is why KEs sharing a term and a direction were treated as one unit in the perturbation analysis (the readout cannot separate them). It should be noted that the genes annotated to a GO BP term capture the toxicologically relevant part of the process a KE describes. GO annotations are assembled from evidence across many biological contexts rather than from toxicology, so a gene set may generally reflect disturbance of a process but miss the crucial elements in a toxicological response.

Mapping the annotations back onto the liver network (Figure 5) shows where they remain absent: the KEs still without a gene set sit predominantly at the organ, individual and population levels, which is precisely where GO offers no terms. To evaluate the impact of our ontology-driven KE-gene set mapping, we analysed pesticide-induced liver toxicity. Pesticides were selected as the focus because they represent a significant dietary exposure route [23], and the liver serves as the primary detoxification organ through intensive pesticide-hepatocyte interactions [24]. Chronic or high-level exposure to pesticides is associated with progressive liver damage, ranging from inflammation (hepatitis) [24] and fibrosis [33] to liver cancer [34], making this compound class highly relevant for the mechanistic evaluation of AOP. We quantified KE perturbation patterns across multiple dose-time conditions for each pesticide using transcriptomics-based enrichment analysis. The four selected compounds act on different molecular targets in the liver. Cycloheximide binds the E site of the 60S ribosomal subunit and stops translation elongation [35]. In rat liver, the resulting loss of protein synthesis rapidly induces hepatocyte apoptosis [36]. Anti-apoptotic proteins with short half-lives are no longer replaced once translation stops, and the block also activates a stress response through CHOP, ATF3 and ATF4 [37, 38]. 2,4-Dinitrophenol is a protonophore. It carries protons back across the inner mitochondrial membrane and removes the gradient that ATP synthase depends on, so hepatocyte respiration continues while ATP synthesis does not [39]. The energy is released as heat, and the liver oxidises more fat to compensate, which is why the compound was sold as a weight-loss agent [39] and why liver-targeted uncouplers are being developed for fatty liver disease [40]. In rats, repeated exposure increases liver weight and produces centrilobular hypertrophy, necrotic foci, and mitochondrial changes [41]. Hexachlorobenzene accumulates in liver tissue, where it induces microsomal enzymes and lowers uroporphyrinogen decarboxylase activity. Heme synthesis is disrupted, porphyrins build up, and hepatic porphyria follows; this was the cause of the mass poisoning in south-eastern Turkey in the late 1950s. Repeated exposure also enlarges the liver and produces liver tumours in rats [42]. Rotenone binds complex I of the mitochondrial electron transport chain and prevents electron transfer. ATP production in hepatocytes falls, and electrons leaking to oxygen form reactive oxygen species [43]. In rats, rotenone raises serum ALT and AST, damages mitochondria and endoplasmic reticulum, and shifts the balance of pro- and anti-apoptotic proteins towards hepatocyte apoptosis [44].

Although ssGSEA identified significantly perturbed KEs based on KE-associated gene sets, these findings should be interpreted as evidence of molecular perturbation rather than direct proof of KE activation or causal progression through an AOP. Transcriptomic changes do not necessarily translate into functional changes at the protein, cellular, or tissue levels [45]. Furthermore, establishing causality within an AOP requires additional evidence, including biological plausibility, key event essentiality, and empirical support for key event relationships [46].

The two panels of Figure 6 describe different aspects of the response. The UpSet plot (Figure 6(B)) records whether a KE was perturbed, while the principal component analysis (Figure 6(A)) uses the scores themselves, so it also reflects how strongly and in which direction each KE responded. This is why the compounds can share most of their perturbed KEs and still be separated in the ordination. Cycloheximide is the clearest example. It shares the majority of its perturbed KEs with the other compounds, yet its scores are markedly more negative (mean NES -1.18, against -0.28 to +0.06 for the others), and it accounts for 214 of the significant negative calls. On the KEs that define PC1, all of which concern lipid and xenobiotic metabolism, cycloheximide reaches values between -1.6 and -3.1 while the other compounds remain close to zero. Its separation therefore does not come from perturbing different KEs, but from perturbing the same KEs far more strongly and almost uniformly downwards, which is consistent with a block of translation elongation suppressing biosynthetic and metabolic programmes across the board. The remaining three compounds were not separated by more than the spread of their own conditions, so their apparent differences in Figure 6(A) should not be interpreted as compound effects. The wider spread of hexachlorobenzene reflects in part its nine conditions against six for the others.

The 20 KE units that responded to every compound (Table 4) cover mitochondrial energy metabolism, lipid handling, DNA damage, inflammation and tissue remodelling, which are the processes a hepatocyte engages under most forms of chemical stress. They therefore describe a general response of the liver network rather than a signature of any one compound. Mitochondrial function and energy metabolism is the largest category, with 8 of the 20 KE units. It includes the complex I events NADH dehydrogenase (ubiquinone) activity (KE 887) and mitochondrial electron transport, NADH to ubiquinone (KE 888), which are the direct target of rotenone, together with oxidative phosphorylation (KE 1545) and proton motive force-driven ATP synthesis (KE 1544; KE 1655), on which 2,4-dinitrophenol acts as an uncoupler. That cycloheximide and hexachlorobenzene perturb the same events is less obvious, since neither targets mitochondria. Blocking translation stops the supply of new respiratory chain subunits, and disturbed heme synthesis limits the cytochromes these complexes require. All four compounds therefore converge on mitochondrial KEs, by different routes. The lipid category shows that the arm of the network leading to steatosis responds to all four exposures, through fatty acid transport (KE 54; KE 465), cholesterol homeostasis (KE 1289) and very-low-density lipoprotein particle assembly. A shared perturbation here does not imply a shared direction of change: 2,4-dinitrophenol increases hepatic fat oxidation and lowers triglyceride content, while rotenone accumulates lipid droplets in rat liver. A KE was counted as perturbed based on the magnitude of its score rather than its sign, so compounds moving the same process in opposite directions both contribute to the shared set. Two processes appear twice in Table 4. Very-low-density lipoprotein particle assembly appears as KE 879; KE 880 and as KE 450, and angiotensin maturation as KE 1743 and KE 1740. In each case the KEs carry the same GO term but differ in direction, so they share one gene set and necessarily receive the same score. None of the four compounds is a classical genotoxicant, and the gene sets for DNA repair (KE 493) and signal transduction in response to DNA damage (KE 1194) overlap considerably with general transcriptional stress programmes, so their perturbation is more likely to reflect a broad stress response than direct DNA damage. Our findings align with previous studies reporting mitochondrial function and energy metabolism [47, 48], lipid metabolism and transport [49, 50], DNA damage response and cell death [51, 52, 53], inflammation and immune signalling [54], and tissue remodelling and fibrosis [55] as hallmarks of hepatotoxicity.

The analysis covered four pesticides and a single human *in vitro* transcriptomics dataset, so the perturbation patterns reported here require testing against a broader range of chemicals and independent datasets. Each condition in Open TG-GATEs is represented by only two biological replicates, which is low for differential expression analysis and makes the p-values entering the ranking metric less stable. Scoring gene sets rather than individual genes limits the effect, but a design with more replicates would give more reliable perturbation scores. The exposure design also bounds what can be concluded. In Open TG-GATEs, the highest concentration of each compound is set from a cell viability assay, so cells are exposed at a level producing only around 10% cytotoxicity [56]. The fiftyfold spread in concentration between compounds therefore reflects differences in potency rather than differences in exposure severity, and the responses reported here are those of largely viable hepatocytes. This argues against the shared perturbations being simple cell death, but it also means that events requiring higher exposure or longer duration are not captured. Time coverage is also uneven as only hexachlorobenzene was measured at 2 h, so early responses cannot be compared across compounds. Gene set size thresholds excluded 50 of the 151 annotated liver KEs from scoring. Of these, 41 had fewer than 10 genes, which may yield unstable enrichment estimates. The remaining 9 had more than 800 genes, representing overly broad processes that hinder mechanistic interpretation. A third of the annotated network was therefore not assessed, and the excluded small sets are those carrying the most specific annotations. Lowering the lower threshold would admit them at the cost of less stable enrichment estimates. Finally, this work makes individual KEs assessable but does not quantify KERs or predict adverse outcomes. Quantifying causality between sequential KEs requires relating how much an upstream KE changes to how much the KE it leads to changes, across a dose range, which needs mathematical modelling on top of the measurements. Forecasting toxicological endpoints additionally requires mechanistic validation, representing important directions for future work.

This work provides a route from a KE to a gene set, and therefore from an AOP to a measurement. The annotations are available on Zenodo [30] and can be reused directly or returned to AOP-Wiki, where they would support any analysis that depends on linking KEs to molecular data.

## 5 Conclusion

Most KEs in AOP-Wiki cannot be measured with transcriptomics, because they carry no Biological Process annotation from which a gene set can be derived. We showed that this gap can be closed systematically. Annotating KEs with fitting GO BP terms raised the proportion of transcriptomics-linkable KEs across the brain, liver, lung and kidney networks, and revising the annotations already present made the resulting gene sets match the events they describe more closely. In a liver case study, most of the KEs perturbed by four pesticides carried annotations added or revised in this work, and the perturbed events included complex I and mitochondrial ATP synthesis, consistent with the steps rotenone inhibits and 2,4-dinitrophenol uncouples. If these findings hold for a wider range of chemicals, a single transcriptomics experiment can report which KEs a compound perturbs across an organ network. Transcriptomics data will then serve as mechanistic evidence in chemical safety assessment, rather than only as a list of changed genes.

## Supporting information

SPARQL queries

AOP network information

Revising the pre-annotated GO terms

All revisions made to the pre-existing liver annotations

## Supplementary information

All SPARQL queries used in this study are provided in Supplementary Files S1. Supplementary Table 1 lists the five hub KEs of each organ network with their degree and betweenness centrality. Supplementary Table 2 lists the obsolete GO terms replaced by their current equivalents and the direction-carrying terms replaced by direction-neutral ones across the four networks. Supplementary Table 3 lists all revisions made to the pre-existing liver annotations, giving the original and replacement term for each KE.

## Acknowledgements

This study has received funding from the European Food Safety Authority (EFSA) under the call reference GP/EFSA/ED/2022/01.

## Declarations

### 6 Author contributions

S.A. performed the curation, analysis and visualisation, and wrote the manuscript. M.M. supervised the work and reviewed the annotations. All authors contributed to the study design, revised the manuscript and approved the final version.

### 7 Data Availability

The GO Biological Process annotations generated in this study for the brain, kidney, liver, and lung AOP networks are publicly available on Zenodo [30]. All scripts developed for data retrieval, annotation, network construction, and transcriptomics analysis are available on GitHub at https://github.com/SidraAdil/GO2KE.

